# Genomic analyses identify key molecules and significant signaling pathways in AZIN1 regulated prostate cancer cells

**DOI:** 10.1101/2022.01.29.478331

**Authors:** Tingxiang Chang, Hanming Gu, James Liu

## Abstract

Antizyme inhibitor 1 (AZIN1) is a critical target in prostate cancer, which regulates the adenosine-to-inosine (A to I) RNA editing during the cancer progression. However, the potential signaling pathways and functions remain unknown. Here, our objective is to figure out the functional molecules and signaling pathways by analyzing the RNA-seq data. The GSE189379 was produced by the Illumina HiSeq 2000 (Homo sapiens). The KEGG and GO analyses showed that focal adhesion and proteoglycans are the mainly affected processes in prostate cancer with the loss of AZIN1. Moreover, we identified ten key molecules including FN1, HRAS, CCND1, RAD51, PCNA, TYMS, CASP3, RRM2, BIRC5, and CCNE2. Therefore, this study provides novel knowledge of AZIN1 mediated prostate cancer.

## Introduction

Prostate cancer is the second most common cancer in men worldwide, and the incidence has elevated over ten years^1^. Prostate cancer is not common under the age of 40, but the incidence is inclined with age^2^. Studies showed environmental factors are critical for the development of prostate cancer^1^. Environmental factors include the high intake of saturated fat and low dietary selenium, vitamin E, and vitamin D^3^. Reports showed that less than 5% of prostate cancer is hereditary, and the major reason for prostate cancer is caused by genetic mutational events^1^. Deficient tumor-suppressive genes such as p53 and p73 appear to be the main cause for the development of prostate cancers^4^. However, the molecular mechanism of prostate cancer is still unclear. Antizyme inhibitor 1 (AZIN1) plays an important role in the polyamine biosynthetic pathway, which is a potential target for cancers^5^. AZIN1 was first found in the liver as an antizyme inhibitor (AZI), which contains two isoforms and involves tumorigenesis and aggressive behaviors^6^. AZIN1 gene is related to the survival and accelerated distant metastases in breast cancers, and the expression of AZIN1 is highly expressed in stomach, lung, and prostate cancers^7^. Overexpression of AZINJ1 promotes tumor growth by maintaining polyamine homeostasis and cell proliferation^8^. Moreover, A to I RNA editing of AZIN contributes to cancer through regulating the polyamine^5^.

In this study, we explored the effects of AZIN1 on prostate cancer by using the RNA-seq data. We identified several DEGs and significant pathways through the KEGG and GO. We also construct the gene enrichment map and protein-protein interaction (PPI) network to study the relationships among the DEGs. These DEGs and biological processes will provide guidance on the treatment of prostate cancers.

## Methods

### Data resources

Gene dataset GSE189379 was downloaded from the GEO database. The data was produced by the Illumina HiSeq 2000 (Homo sapiens) (KU Leuven, Herestraat 49, Leuven, Belgium). The analyzed dataset includes 3 of controls and 3 of AZIN1 siRNA treated PC3 cells.

### Data acquisition and processing

The data were organized and conducted by R package as previously described^9–12^. We used a classical t-test to identify DEGs with P< 0.05 and fold change ≥1.5 as being statistically significant.

The Kyoto Encyclopedia of Genes and Genomes (KEGG) and Gene Ontology (GO) KEGG and GO analyses were conducted by the R package (ClusterProfiler) and Reactome. P<0.05 was considered statistically significant.

### Protein-protein interaction (PPI) networks

The Molecular Complex Detection (MCODE) was used to create the PPI networks. The significant modules were produced from constructed PPI networks. The pathway analyses were performed by using Reactome (https://reactome.org/), and P<0.05 was considered significant.

## Results

### Identification of DEGs in prostate cancer cells with deficient AZIN1

To determine the impact of AZIN1 on prostate cancer cells, we analyzed the RNA-seq data from the PC3 cells (prostate cells) with the knockout of AZIN1. A total of 1172 genes were identified with the threshold of P < 0.05. The up-and-down-regulated genes were shown by the heatmap and volcano plot (Figure 1). The top ten DEGs were selected in Table 1.

**Figure 1.**
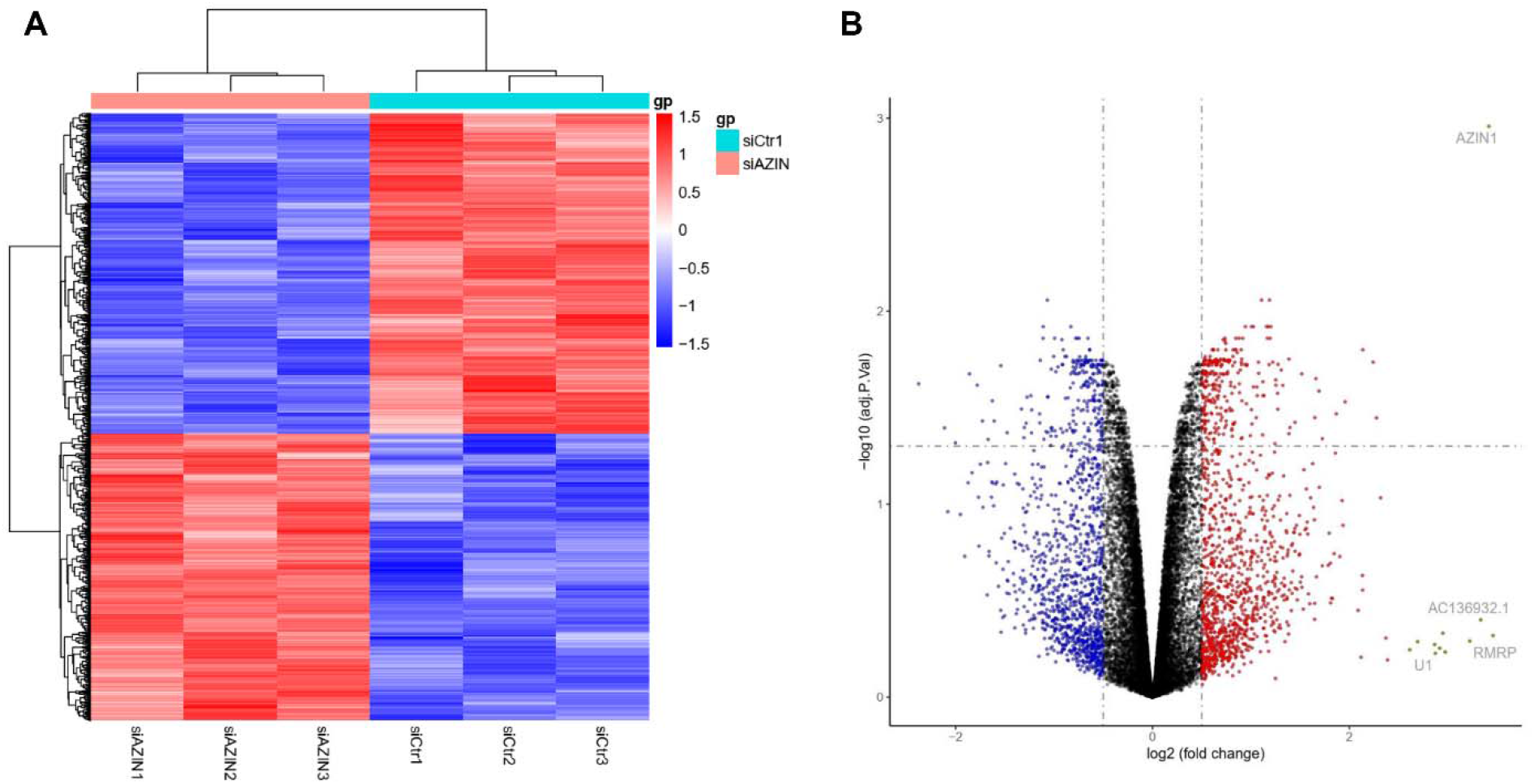
Heatmap and volcano plot in prostate cancer cells with deficient AZIN1. (A) Heatmap of significant DEGs in prostate cancer cells with deficient AZIN1. Significant DEGs (P < 0.05) were used to create the heatmap. (B) Volcano plot for DEGs in prostate cancer cells with deficient AZIN1. The most significantly changed genes are highlighted by grey dots.

**Table 1.**
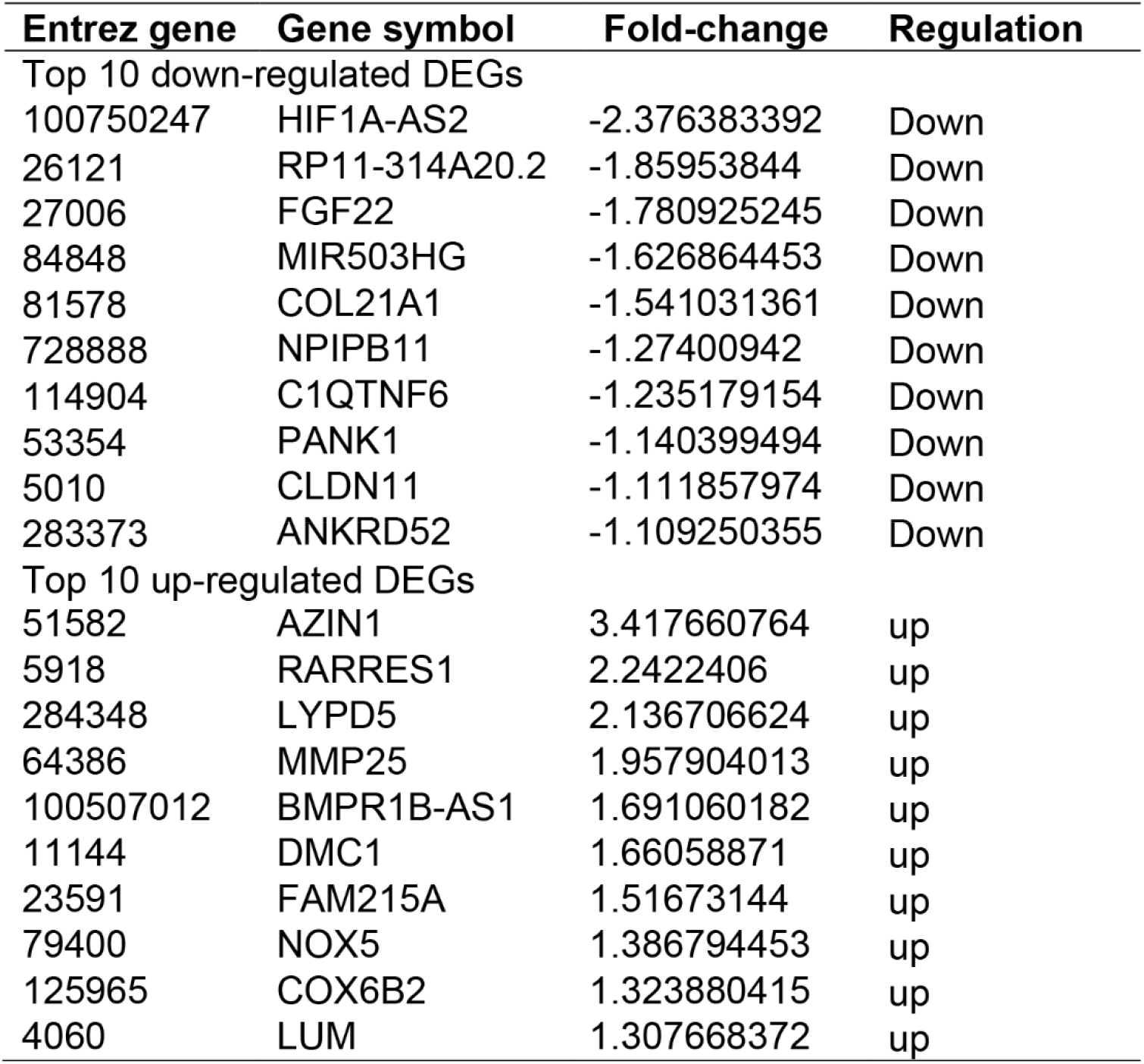

### Enrichment analysis of DEGs by KEGG and GO analyses

To further figure out the mechanism of AZIN1 regulated cancer cells, we performed the KEGG and GO analyses (Figure 2). We identified the top ten KEGG signaling pathways including “Focal adhesion”, “Proteoglycans in cancer”, “Gastric cancer”, “Hepatocellular carcinoma”, “Cell cycle”, “Small cell lung cancer”, “Cellular senescence”, “Colorectal cancer”, “Prostate cancer”, and “Pancreatic cancer”. We also identified the top ten biological processes of GO including “Reproductive structure development”, “Reproductive system development”, “cell–substrate adhesion”, “DNA replication”, “sex differentiation”, “cell-matrix adhesion”, “gonad development”, “DNA–dependent DNA replication”, “male sex differentiation”, and “regulation of cell–matrix adhesion”. We identified the top ten cellular components of GO including “collagen–containing extracellular matrix”, “focal adhesion”, “cell–substrate junction”, “endoplasmic reticulum lumen”, “basement membrane”, “protein kinase complex”, “serine/threonine protein kinase complex”, “cyclin-dependent protein kinase holoenzyme complex”, “DNA replication preinitiation complex”, and “laminin complex”. We identified the top ten molecular functions of GO including “phosphoric ester hydrolase activity”, “small GTPase binding”, “GTPase binding”, “extracellular matrix structural constituent”, “phosphoprotein phosphatase activity”, “protein tyrosine phosphatase activity”, “cyclin-dependent protein serine/threonine kinase regulator activity”, “2-oxoglutarate-dependent dioxygenase activity”, “single-stranded DNA helicase activity”, and “1-phosphatidylinositol binding”.

**Figure 2.**
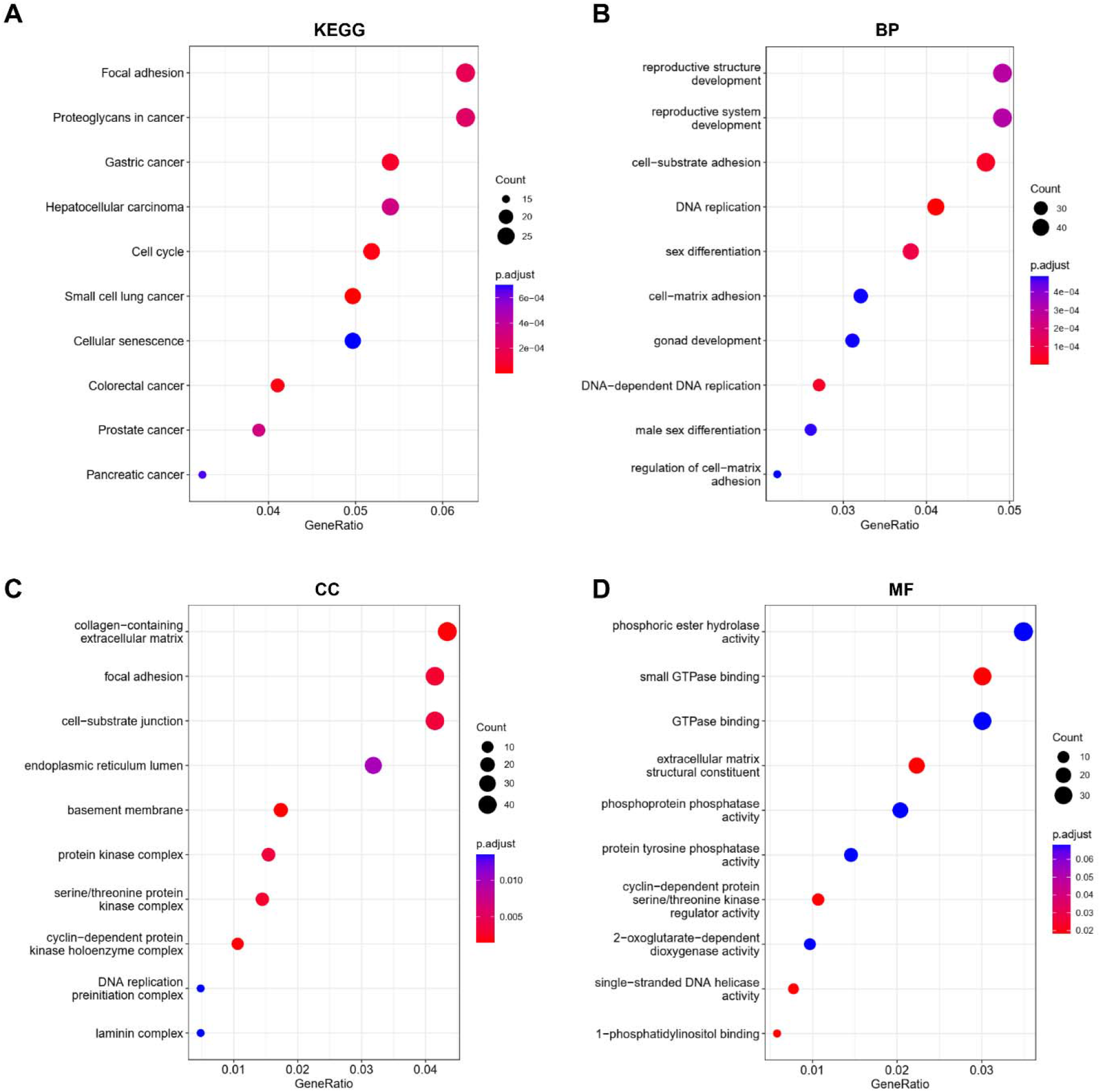
KEGG and GO analyses of DEGs in prostate cancer cells with deficient AZIN1. (A) KEGG analysis, (B) Biological processes, (C) Cellular components, (D) Molecular functions.

### PPI networks in prostate cancer cells with deficient AZIN1

To determine the relationship among the DEGs, we constructed the PPI networks by using the Cytoscope software (combine score > 0.4). Table 2 showed the top ten interactive molecules with the highest degree scores. The top two modules were indicated in Figure 3. We further explored the mechanisms with Reactome map (Figure 4) and identified the top ten significant processes including “G1/S-Specific Transcription”, “Mitotic G1 phase and G1/S transition”, “G1/S Transition”, “NR1H3 & NR1H2 regulate gene expression linked to cholesterol transport and efflux”, “NR1H2 and NR1H3-mediated signaling”, “Downstream signaling of activated FGFR1”, “Crosslinking of collagen fibrils”, “Assembly of collagen fibrils and other multimeric structures”, “Extracellular matrix organization”, and “PI-3K cascade:FGFR1” (Supplemental Table S1).

**Figure 3.**
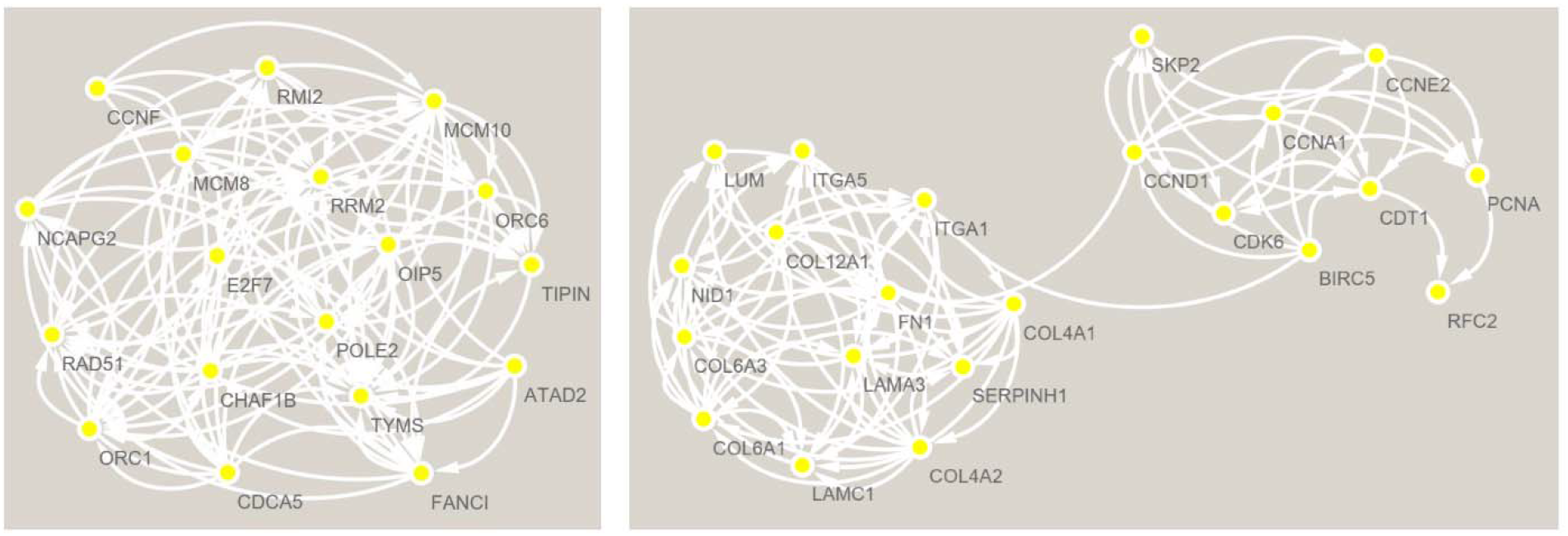
The PPI network analyses of DEGs in prostate cancer cells with deficient AZIN1. The top two clusters were constructed by MCODE according to the DEGs.

**Figure 4.**
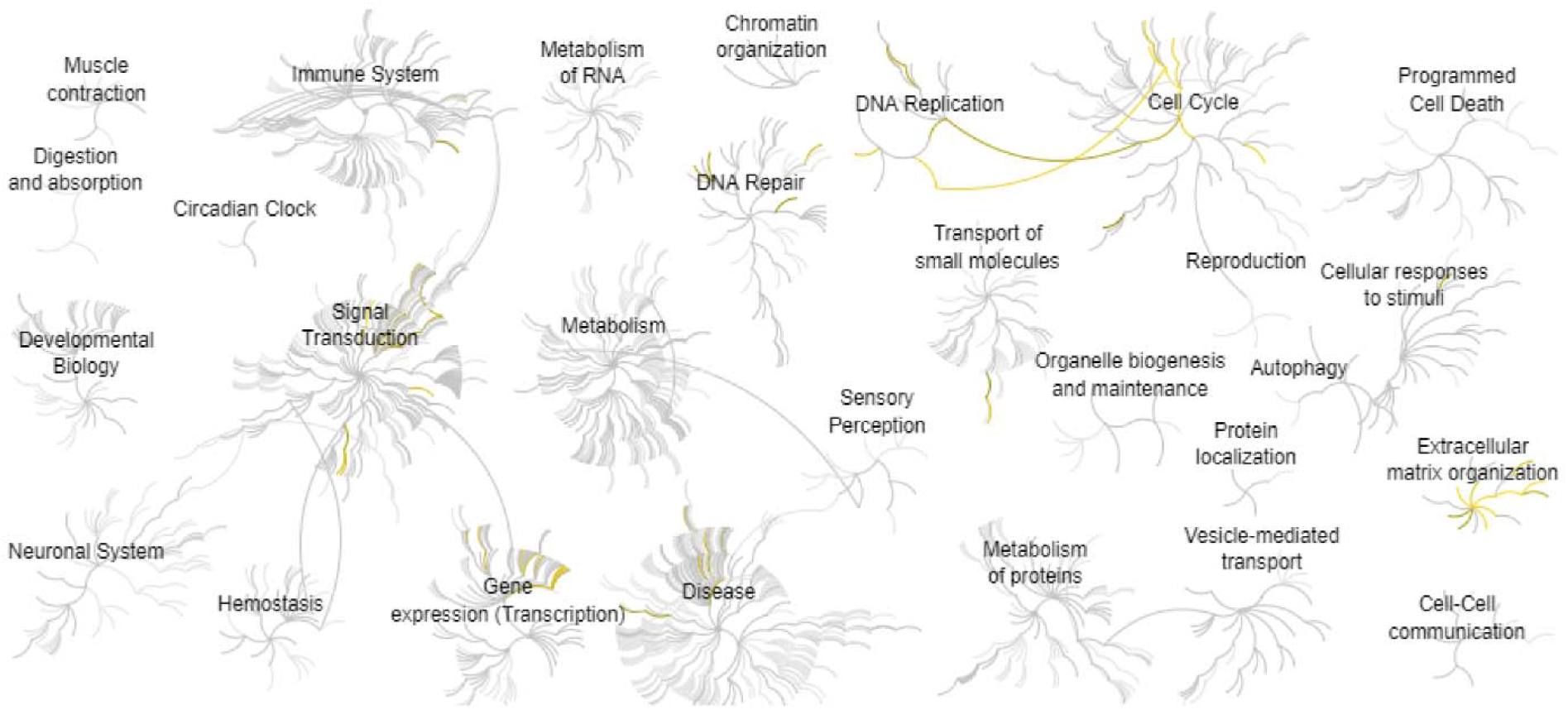
Reactome map representation of the significant biological processes in prostate cancer cells with deficient AZIN1.

**Table 2.** Top ten genes demonstrated by connectivity degree in the PPI network.

| **Gene symbol** | **Gene title** | **Degree** |
| --- | --- | --- |
| FN1 | Fibronectin 1 | 53 |
| HRAS | Hras Proto-Oncogene, Gtpase | 48 |
| CCND1 | Cyclin D1 | 43 |
| RAD51 | RAD51 Recombinase | 33 |
| PCNA | Proliferating Cell Nuclear Antigen | 32 |
| TYMS | Thymidylate Synthetase | 31 |
| CASP3 | Caspase 3 | 30 |
| RRM2 | Ribonucleotide Reductase Regulatory Subunit M2 | 29 |
| BIRC5 | Baculoviral IAP Repeat Containing 5 | 27 |
| CCNE2 | Cyclin E2 | 27 |

## Discussion

AZIN1 is one of the most frequently occurring A-to-I RNA alterations in cancers and the overexpression of AZIN1 is associated with aggressive tumors and elevated ornithine decarboxylase and polyamines accumulation^7^. However, the molecular function of AZIN1 in prostate cancer remains unexplored.

Our study showed that the deficient AZIN1 in prostate cancer cells mainly affects the focal adhesion and proteoglycans processes. The study by Sheila Figel et al found that focal adhesion kinase is highly expressed in prostate cancer, which enhances the growth, survival, migration, metastasis of prostate tumors^13^. Thomas R Johnson et al also found focal adhesion kinase regulates the aggressive phenotype of prostate cancer cells^14^. Proteoglycans regulate cell signaling, adhesion, growth, and apoptosis, and the change of proteoglycans is observed in prostate cancers. Proteoglycans may become a tumor regulator through mediating the sonic hedgehog signaling. Moreover, changes in tumors can alter proteoglycan contents and structure to change their functions^15^.

By analyzing the Reactome map and PPI network, we identified the top ten interactive molecules that may involve the process of AZIN1 affected prostate cancers. The study by Dibash K Das et al showed that overexpression of FN1 is positively correlate with aggressive prostate cancer^16^. Clinton L Cario et al found that HRAS is a driver gene in the prostate cancer patients by using the machine learning methods^17^. G protein-coupled receptors (GPCRs) promote cells to respond to several stimuli through generation of second messengers, which in turn regulate a variety of processes such as cell survival, proliferation, migration, and differentiation^18–28^. As a key GTPase, HRAS was found to be involved in controlling the GPCR/cAMP signaling pathways, thereby mediating the cancer cells^29^. Mingming Wang et al found the microRNA-4873-3p inhibits the progression of prostate cancer by targeting CCND1^30^. Maria Nowacka-Zawisza et al proved that the RAD51 polymorphisms is closely related to the high risk of prostate cancer^31^. Huajun Zhao et al found the tyrosine phosphorylation of PCNA is a critical prostate cancer target^32^. Circadian clocks regulate numerous downstream genes and signaling pathways^33–43^. Most of genes indicate the rhythm during the day and night. Interestingly, the oscillation of PCNA was observed during the light and dark cycles, which may affect the proliferation of tumor cells^44^. Christoph Burdelski et al found that the increased thymidylate synthase is related to aggressive tumor features in prostate cancer^45^. As a key apoptosis gene, the CASP3 is upregulated in prostate cancer cell lines^46^. Ying Z Mazzu et al found the RRM2 is increased in prostate cancer, which is a novel mechanism of poor-prognosis^47^. Daixing Hu et al found the gene BIRC5 can be used to construct the prognostic model of prostate cancers^48^. Zhong Wu et al found the cyclin E2 isa PTEN-regulated gene that is related to human prostate cancer metastasis^49^.

In conclusion, our study found a strong relationship between AZIN1 and prostate cancer. The focal adhesion and proteoglycans are the major affected processes during the development of prostate cancer. Our study provides valuable insights for the treatment of prostate cancer.

## Author Contributions

Tingxiang Chang: Methodology and Writing. Hanming Gu, James Liu: Conceptualization, Writing-Reviewing and Editing.

## Funding

This work was not supported by any funding.

## Declarations of interest

There is no conflict of interest to declare.

## Notes

### Competing Interest Statement

The authors have declared no competing interest.

